# Masitinib is an oral, brain penetrant inhibitor of microglial and mast cell activity with neuroprotective potential in progressive forms of multiple sclerosis

**DOI:** 10.64898/2026.07.02.735783

**Authors:** Patrick Vermersch, Alain Moussy, Colin D Mansfield, Olivier Hermine

**Author notes:** Corresponding authors: Olivier Hermine, MD. Imagine Institute, 24 Boulevard Montparnasse, 75014, Paris, France. Colin Mansfield, PhD. AB Science, 3 Avenue George V, 75008, Paris, France. Electronic address. CONTRIBUTORS: Conceptualization: PV, AM, OH; Data interpretation: AM, OH; writing — original draft preparation: CDM; writing — draft review and editing: PV, AM, OH, and CDM. All authors critically reviewed the manuscript and approved the final version for submission. DISCLAIMER: Masitinib is under clinical development by the study funder AB Science. The sponsor was involved in the study design, interpretation of the data, writing of the report, and decision to submit the manuscript for publication.

## Abstract

**Introduction:** Progressive multiple sclerosis (MS), including primary progressive MS (PPMS) and non-active secondary progressive MS (nSPMS), remains an unmet need, as few treatments target innate immune pathways. Masitinib (AB1010) is a selective tyrosine kinase inhibitor that targets c-Kit and colony-stimulating factor 1 receptor pathways. This mechanism disrupts mast cell-microglia interactions, key innate immune effectors in progressive MS pathogenesis, reducing neuroinflammation and neuronal damage. In the phase 3 AB07002 trial, masitinib (4.5 mg/kg/d) over 96 weeks met its primary endpoint. Comparable signals in PPMS and nSPMS indicated masitinib benefited both phenotypes. Secondary analyses showed that masitinib lowered the progression to wheelchair dependence (EDSS ≥7.0, 12 weeks) and reduced the 12-week confirmed EDSS progression risk by 37% versus placebo.

**Methods:** This study aimed to confirm that oral masitinib achieves central nervous system (CNS) concentrations sufficient to modulate CSF1R and wild-type c-Kit, thereby underpinning its neuroprotective potential. Male Sprague Dawley rats (n=12, ∼200 g) were administered a single oral dose (30 mg/kg). Plasma and brain samples were collected at 2, 4, 8, and 24 hours post-dose (n=3 per time point). Masitinib (AB1010) and its metabolite (AB3280) were quantified in plasma and brain homogenates using LC-MS/MS.

**Results:** Masitinib reached a brain Cmax of 223.5 ng/mL (∼450 nM), exceeding IC_50_ values for CSF1R and wild-type c-KIT by ∼5-fold and 2-fold, respectively, indicating effective CNS target engagement. The active metabolite AB3280 also achieved brain Cmax levels with full inhibitory activity. Masitinib demonstrated consistent CNS penetration supported by a proportional plasma-to-brain exposure relationship.

**Conclusion:** Masitinib’s favorable CNS penetration and safety profile, alongside its unique mast cell inhibition, position it as a compelling candidate for progressive MS treatment, either as monotherapy or in combination with other agents. This multifaceted immunomodulatory approach addresses critical unmet needs in progressive MS and supports further clinical development.

## Introduction

### Progressive Multiple Sclerosis: A Critical Unmet Medical Need

Progressive forms of multiple sclerosis (MS), defined as primary progressive MS (PPMS) and non-active secondary progressive MS (nSPMS), are characterized by a steady and continuous worsening of neurological function without distinct relapses or remissions. Globally, an estimated 2.8 million people live with MS, equating to 35.9 per 100,000 individuals [Walton 2020]. Of these, approximately 37% have progressive forms of MS, with 15% having PPMS [Montalban 2017; Miller 2007] and 22% having nSPMS [Chisari 2024; Forsberg 2023]. Unlike relapsing forms of MS, such as relapse-remitting MS (RRMS) and active SPMS, which are partly driven by peripheral adaptive immune system activity and marked by episodes of inflammatory demyelination, progressive MS is driven to a far greater extent by the dysfunctional activity of the innate immune system compartmentalized within the central nervous system (CNS) [Stys 2019; Hendriksen 2017; Fani Maleki 2019; Skaper 2018; Skaper 2014]. This distinct pathophysiology explains the limited efficacy of most current MS therapies in treating progressive MS, as they primarily target adaptive immunity (B and T cells) and fail to adequately address neurodegenerative and innate immune-driven mechanisms.

Current pharmacological treatments for progressive MS are limited in terms of their number and effectiveness. Ocrelizumab, a humanized anti-CD20 monoclonal antibody, is the only approved therapy for PPMS, authorized for early stage disease with inflammatory activity. However, its clinical benefits are modest, offering only a slight reduction in disability progression without any demonstrated improvement in the patient’s quality of life. Currently, no therapies have been approved for nSPMS. Numerous Bruton’s tyrosine kinase (BTK) inhibitors, which target B cell and microglial activation, are in the late phase of clinical development for MS [Vermersch 2025]. Despite some success, development challenges, including safety concerns, have somewhat tempered enthusiasm, and there is an urgent need for additional drugs. This need stems from the complex and multifaceted nature of progressive MS pathophysiology, which involves innate immune mechanisms within the CNS that are not fully addressed by current treatments. Therefore, the development of new therapies capable of penetrating the CNS with more effective modulation of innate immune activity is critical to slow or halt disease progression and meet the substantial unmet medical needs of patients with PPMS and nSPMS.

### The Mechanism of Action of Masitinib in Progressive MS

Masitinib (AB1010) is a selective oral tyrosine kinase inhibitor that targets key innate immune cells involved in the pathophysiology of progressive MS, specifically mast cells and microglia. By inhibiting the colony-stimulating factor 1 receptor (CSF1R), masitinib modulates microglial proliferation and activity, which are critical contributors to neuroinflammation and demyelination in patients with MS. Simultaneously, by targeting c-Kit, Lyn, and Fyn, masitinib regulates mast cell functions, including survival, proliferation, and degranulation, thereby reducing their role in orchestrating inflammatory cascades and disrupting the blood-brain barrier (BBB). Mast cells and microglia interact within the CNS (mast cell-microglia crosstalk) to amplify neuroinflammatory responses, contributing to neuronal damage and disease progression. Masitinib’s dual action on these innate immune cells aims to shift the neuroimmune environment from a neurotoxic to a neuroprotective state by remodeling cellular interactions and reducing chronic inflammation. This mechanism sets masitinib apart as an innovative therapeutic strategy for progressive MS, concentrating solely on modulating innate immunity without affecting adaptive immunity.

### Masitinib Trial AB07002 in Patients with Progressive Forms of MS Met its Primary Endpoint

Study AB07002 (NCT01433497) was a randomized, double-blind, placebo-controlled phase 3 trial comparing masitinib at a dosage of 4.5 mg/kg/d against a placebo in patients with progressive MS who were progressing but not clinically active [Vermersch 2022]. The primary endpoint was the change in the Expanded Disability Status Scale (EDSS) score over 96 weeks, measured using a repeated-measures generalized estimating equation approach. Results were expressed as least-squares mean (LSM) change on the EDSS from baseline (δEDSS, wherein a positive value indicates disability progression), with treatment effect (masitinib vs placebo) reported as the between group difference (ΔLSM, wherein a negative value favors masitinib). This innovative methodology integrates all available data points to achieve a population-averaged interpretation while reducing the sample size required for a given study power compared with the conventional time to confirmed disability progression endpoint. However, its lack of precedent in other MS trials and its disconnection from the original EDSS scores make direct clinical interpretation challenging.

The study was successful in its primary endpoint, with the masitinib 4.5 mg/kg/d group (n=199) showing significant benefit over placebo (n=101) with a δEDSS of 0.001 versus 0.098 (p=0.0256); that is, a ΔLSM of -0.097 (97% CI -0.192 to -0.002) [Vermersch 2022]. This finding was confirmed by multiple sensitivity analyses. On secondary measures, masitinib significantly lowered the risk of progression to wheelchair use (EDSS ≥7.0, 12-week confirmed), with 0% of masitinib-treated patients reaching this milestone versus 4% in the placebo group (p=0.013). Additionally, masitinib reduced the risk of time to first EDSS progression (unconfirmed) by 42% relative to placebo (hazard ratio 0.58, 95% CI, 0.35 to 0.96; p=0.034), and the risk of 12-week confirmed EDSS progression by 37% relative to placebo (hazard ratio 0.63, 95% CI, 0.33 to 1.20; p=0.159), although the study was underpowered for these endpoints [Vermersch 2022]. In comparison, ocrelizumab in PPMS and tolebrutinib in nSPMS had relative risk reductions of 12-week confirmed EDSS progression of 24% (p=0.03) and 24% (p=0.013), respectively [Montalban 2017; Fox 2025].

Subgroup analysis showed comparable signals in patients with PPMS and nSPMS, indicating that masitinib could benefit both disease phenotypes equally, although the study was underpowered for these patient populations. Considering the primary endpoint in the PPMS subgroup, ΔLSM was -0.128 (95% CI -0.285 to -0.028) in masitinib-treated PPMS patients (n=79) versus placebo PPMS patients (n=45) [Vermersch 2022]. The relative risk of 12-week confirmed EDSS progression was reduced by 59% in masitinib-treated patients (hazard ratio 0.41, 95% CI 0.14 to 1.23; p=0.111), and the relative risk of unconfirmed EDSS progression was reduced by 61% (hazard ratio 0.39, 95% CI 0.17 to 0.85; p=0.183). Similarly, for the nSPMS subgroup, ΔLSM was -0.104 (95% CI -0.198 to -0.008) in masitinib-treated nSPMS patients (n=120) versus placebo nSPMS patients (n=56) [Vermersch 2022]. The relative risk of 12-week confirmed EDSS progression was reduced by 26% in masitinib-treated patients (hazard ratio 0.74, 95% CI 0.29 to 1.85; p=0.514) and the relative risk of unconfirmed EDSS progression was reduced by 35% (hazard ratio 0.65, 95% CI 0.32 to 1.36; p=0.254).

Given the critical roles of microglia and mast cells in the pathophysiology of progressive MS, it is desirable to achieve therapeutically relevant CNS concentrations of masitinib (AB1010) and/or its active metabolite AB3280 for direct intracerebral modulation of these cellular targets. Herein, we present preclinical evidence that masitinib can penetrate the BBB at therapeutically relevant concentrations, supporting its potential to engage key immune mechanisms implicated in progressive MS.

## Methods

Twelve adult male Sprague Dawley rats, each weighing approximately 200 g, were administered a single oral dose of masitinib at 30 mg/kg. Animals were housed and maintained under standard laboratory conditions with free access to food and water. Plasma and brain tissue samples were collected at predetermined time points of 2, 4, 8, and 24 hours post-administration, with three rats sacrificed at each time point to obtain biological specimens. Blood samples were collected via cardiac puncture under appropriate anesthesia, and plasma was separated by centrifugation. Brain tissues were rapidly excised, rinsed in ice-cold saline to remove residual blood, and homogenized for analysis.

Quantitative analysis of masitinib and its active metabolite AB3280 in plasma and brain homogenates was performed using a validated liquid chromatography-tandem mass spectrometry (LC-MS/MS) method.

Chromatographic separation was performed using a C18 reverse-phase column under optimized conditions. Calibration curves were constructed using known standards, and quality control samples ensured assay accuracy and precision. Sample preparation involved protein precipitation and extraction protocols suitable for both plasma and brain matrices. All experimental procedures involving animals were conducted in accordance with institutional and national ethical guidelines for animal care and use. Detailed procedures are described in the Supplementary Appendix.

For the in vitro evaluation of masitinib permeability, data were referenced from a porcine brain endothelial cell (PBEC) monolayer model [Di Marco 2019]. This model used primary PBEC cultures to mimic the properties of the BBB. The permeability coefficient of masitinib was inferred by comparing it to that of imatinib, a structurally related tyrosine kinase inhibitor, based on previously reported permeability values obtained under standardized assay conditions.

## Results

The pharmacokinetic data indicates that masitinib (AB1010) and its active metabolite (AB3280) effectively crossed the BBB, achieving therapeutically relevant concentrations in the CNS (see Table 1 and Supplemental Appendix Tabe S1). After a single oral dose of 30 mg/kg, masitinib exhibited a brain maximum concentration (C_max_) of 223.5 ng/mL, equivalent to approximately 450 nM, which exceeded the inhibitory concentration 50% (IC_50_) values for its primary targets, CSF1R and wild-type c-KIT, by substantial margins. Specifically, the brain C_max_ surpassed the CSF1R IC_50_ (90 nM) [Trias 2016] by approximately 5-fold and the c-Kit IC_50_ (200 nM) [Dubreuil 2009] by more than 2-fold, indicating effective target engagement within the brain. Similarly, the active metabolite AB3280 reached a brain C_max_ of 33.8 ng/mL and retained its full inhibitory activity against these targets. The proportional relationship observed between the plasma area under the curve (AUC) and brain drug exposure further supports the consistent CNS penetration of masitinib and its metabolite.

**Table 1:** Pharmacokinetic parameters of AB1010 and AB3280 in male Sprague Dawley rats.

| Group | C max<br>(ng/mL) | T max<br>(h) | AUC t<br>(ng/mL*h) | AUC inf<br>(ng/mL*h) | %AUC<br>extra | t 1/2<br>(h) | AUC brain / AUC<br>plasma |
| --- | --- | --- | --- | --- | --- | --- | --- |
| **AB1010**<br>**(brain)** | 223.51 | 4 | 2485 | 2598 | 4.35 | 5.19 | 23% |
| **AB1010**<br>**(plasma)** | 1492.57 | 4 | 10719 | 10742 | 0.21 | 2.62 | NC |
| **AB3280**<br>**(brain)** | 33.84 | 8 | 626 | NC | NC | NC | 84% |
| **AB3280**<br>**(plasma)** | 117.33 | 4 | 746 | 762 | 2.18 | 4.09 | NC |

AB1010, masitinib; AB3280, active metabolite of masitinib; Cmax, maximum observed concentration; Tmax, time to reach Cmax; AUCt, area under the concentration–time curve from time zero to the last measurable concentration; AUCinf, area under the concentration–time curve from time zero extrapolated to infinity; %AUCextra, percentage of AUCinf attributable to extrapolation; t½, apparent terminal elimination half-life; AUC, area under the concentration–time curve (extent of exposure); AUC brain/AUC plasma (Kp,brain): Ratio of the area under the concentration–time curve in brain tissue to that in plasma; a measure of the extent of drug penetration across the blood–brain barrier.

The in vivo evidence supporting the ability of masitinib to cross the BBB is further bolstered by independent in vitro findings [Di Marco 2019]. Using a porcine brain endothelial cell (PBEC) monolayer model, researchers reported a permeability coefficient for masitinib of 20.4 × 10^-6^ cm/s, which closely aligns with that of imatinib (20.6 × 10^-6^ cm/s). The in vitro permeability threshold was set at 3 × 10□□ cm/s, with values above indicating medium to high permeability. The PBEC model was validated through a correlation analysis involving 36 structurally diverse compounds, including imatinib, with an R^2^ value of 0.73, confirming its predictive capacity. Given the structural similarities between masitinib and imatinib, their comparable permeability values, and the validated in vitro–in vivo correlation framework, masitinib is predicted to exhibit similar in vivo BBB penetration as imatinib (9.3 × 10^-6^ cm/s), that is, moderate permeability.

These findings provide proof of concept that oral administration of masitinib can achieve CNS concentrations sufficient to modulate microglial and mast cell activity. Translating these results to human dosing, masitinib doses of 3.0 to 4.5 mg/kg are expected to yield comparable CNS exposure, considering the additive effects of AB3280. Furthermore, independent binding kinetics analysis has shown that masitinib acts as a type II kinase inhibitor, characterized by slow dissociation kinetics that confer prolonged target engagement [Georgi 2018]. This extended target occupancy suggests sustained inhibition of mast cell and microglia activation beyond what IC_50_ values alone would indicate.

## Discussion

### Innate Immune Dysregulation: Microglia and Mast Cells in Progressive MS Pathogenesis

As evident from the growing body of literature, innate immune activity is at the cutting edge of theories regarding the underlying pathophysiological mechanisms of progressive MS, with a consensus that drugs aimed at these targets have strong therapeutic potential. It is now well established that microglia play a central role in the pathogenesis of progressive MS, significantly contributing to lesion development and sustaining the chronic CNS-compartmentalized inflammation associated with neurodegeneration during the progressive phase of MS [Vermersch 2025; Van den Bosch 2024; Kamma 2022; Muzio 2021]. Microglia exhibit remarkable plasticity, adopting diverse phenotypes that can either support neuroprotection or drive neurotoxicity, depending on their activation state [Mado 2023]. Transcriptomic and proteomic profiling has indicated that microglia are heterogeneous and may shift their state depending on their microenvironment [Mahmood 2022]. A review by Yong highlighted this dualistic nature of microglia in MS, noting their beneficial roles in the early stages of the disease, but observing a shift toward harmful phenotypes as the disease progresses [Yong 2022]. Consequently, therapeutic strategies are increasingly aimed at modulating microglial function to restore homeostasis and steer them away from neurodegenerative states, rather than depleting them. Supporting this approach, the CSF1R signaling pathway has been identified as a crucial target for therapeutic intervention, suggesting that targeting CSF1R could modulate microglial proliferation and molecular identity to mitigate harmful neuroinflammation in progressive MS [Hagan 2020]. Collectively, these findings position microglia as pivotal modulators of the disease trajectory in progressive MS, with their functional state directly influencing the pathological outcomes and therapeutic potential.

Mast cells are important, yet underreported, contributors to progressive MS pathogenesis through interconnected mechanisms that amplify neuroinflammation, disrupt the blood-brain barrier (BBB), and promote demyelination and neurodegeneration [Pinke 2020a; Ribatti 2020; Jones 2019; Skaper 2018]. The potential role of mast cells in MS was clearly demonstrated through the experimental autoimmune encephalomyelitis (EAE) model, an animal model of human demyelinating diseases, where mast cell-deficient mice showed reduced disease severity and reintroduction of mast cells restored full disease [Secor 2000]. Inhibitors of mast cell activity likewise supported the therapeutic premise of targeting mast cells in MS, as treatment of EAE mice with the tyrosine kinase inhibitor masitinib showed a significant, dose-dependent reduction in disease relative to control mice [Vermersch 2012], while the mast cell stabilizer ketotifen fumarate has been shown to trigger a protective effect characterized by lower disease prevalence and severity of EAE and restored CNS-barrier permeability [Pinke 2020b]. Numerous studies support the direct involvement of mast cells in MS pathogenesis, primarily through the release of mediators such as histamine, tryptase, chymase, heparin, serotonin, nitric oxide, proteases, cytokines (IL-6, IL-8, TNF-α), and chemokines, which promote inflammatory cell recruitment, activate T cells, especially Th1 and Th17, and sustain the inflammatory environment of MS [Brown 2018; Hendriksen 2017; Costanza 2012; Christy 2007; Secor 2000; Dines 1997]. The abundance of mast cells is notably higher in the CNS of patients with MS than in healthy controls, especially near blood vessels and demyelinating plaques [Krüger 2018; Couturier 2008; Krüger 2001; Ibrahim 1996]. Mast cells, strategically positioned near blood vessels, disrupt the BBB by releasing vasoactive mediators and cytokines, degrading BBB components, and increasing permeability [Yue 2023; Shelestak 2020; Ribatti 2015; Dong 2014; Sayed 2010]. For example, mast cell-derived TNF-α and chemokines upregulate endothelial adhesion molecules, promoting leukocyte CNS migration and neuroinflammation; histamine’s action on H2 receptors enhances BBB permeability, aiding protein and immune cell passage; and meningeal mast cells release TNF-α, recruiting neutrophils that destabilize the BBB and promote T-cell infiltration. Beyond BBB disruption, mast cells contribute to demyelination and axonal injury by releasing proteases and histamine, which degrade myelin proteins. Myelin breakdown products, such as MBP and P2, can further stimulate mast cell degranulation, creating a feedback loop that worsens tissue damage [Johnson 1988]. Mast cells also interact with Th17 cells and other immune populations, modulating the autoimmune response against myelin and oligodendrocytes [Elieh-Ali-Komi 2017].

Finally, mast cells participate in neuroimmune interactions by communicating with microglia and other glial cells, significantly influencing the CNS microenvironment and implicating themselves in neuroinflammation and neurodegeneration [Jones 2019]. Pro-inflammatory cytokines from mast cells activate microglia, creating a feedback loop that exacerbates neurodegeneration and accelerates disease progression [Skaper 2018; Skaper 2014; Skaper 2012]. It has been proposed that the initiation, propagation, and perpetuation of chronic neuroinflammation depend on mast cell-microglia crosstalk [Sandhu 2021].

Collectively, these findings highlight mast cells as key contributors to the immunopathology of progressive MS, orchestrating the chronic neuroinflammatory environment characteristic of progressive MS through complex interactions with microglia. Furthermore, mast cells actively drive MS pathogenesis through the release of inflammatory mediators, BBB disruption, immune cell recruitment, and direct degradation of myelin. Therefore, targeting mast cell activity may mitigate disease progression and improve outcomes.

### BTK Inhibitors Targeting Microglia in Progressive Multiple Sclerosis: Clinical Trial Insights and Therapeutic Implications

It is now widely recognized that microglia are crucial in the development of progressive MS, playing a significant role in lesion formation and maintaining chronic inflammation within the CNS, which is linked to neurodegeneration during the progressive stage of MS. Perhaps the most compelling evidence to date is the successful pharmacological targeting of microglia in the tolebrutinib HERCULES phase 3 study involving individuals with nSPMS [Fox 2025]. Tolebrutinib is a drug that targets disease-associated microglia and B cells within the CNS by inhibiting BTK. These findings were further supported by a pooled analysis of data from two other studies, GEMINI 1 and 2, which indicated that tolebrutinib treatment in relapsing MS delayed disability accumulation, likely by addressing microglia-associated smoldering compartmentalized inflammation within the CNS [Oh 2025].

Thus far, the BTK inhibitors evobrutinib, tolebrutinib, and fenebrutinib have been clinically evaluated in MS populations with mixed results. The tolebrutinib HERCULES trial (ClinicalTrials.gov Identifier: NCT04411641) achieved its primary endpoint by demonstrating delayed confirmed disability progression in patients with nSPMS [Fox 2025]. Conversely, the tolebrutinib PERSEUS trial (NCT04458051) in patients with PPMS did not meet its primary endpoint [Sanofi Press Release 2025]. The FENtrepid trial (NCT04544449) found that fenebrutinib was at least as effective as ocrelizumab (non-inferior study design) in slowing disability progression in patients with PPMS [Roche Press Release 2025], while the FENhance trials of fenebrutinib versus teriflunomide in relapsing MS (NCT04586010 and NCT04586023) also met their primary endpoints (reduced annualized relapse rate over 96 weeks) but showed no significant reduction in the risk of 12-week composite confirmed disability progression. Furthermore, an imbalance with respect to reported fatalities across fenebrutinib treatment arms was observed, raising additional safety concerns [Roche Press Release 2026]. Finally, evobrutinib failed to demonstrate efficacy in two relapsing MS trials (NCT04338022 and NCT0433806) [Montalban 2024],

### Comparative Discussion of Masitinib and BTK Inhibitors in Progressive Multiple Sclerosis

Masitinib also targets microglia by inhibiting the CSF1R signaling pathway, with an important feature being that it inhibits excessive microglial proliferation and the emergence of aberrant glial cells without resulting in the depletion of all microglial cells [Trias 2016]. Furthermore, unlike BTK inhibitors, masitinib does not target B-cell activity and therefore does not suppress immune function. This is particularly relevant for patient populations that require long-term treatment and who, in certain cases, already have a weakened immune system due to previous treatments or because of their age. The results of study AB07002 showed that there was no evidence of an elevated risk of infection for masitinib relative to placebo over the 96-week treatment period in this population [Vermersch 2022]. The safety profile of masitinib in study AB07002 was consistent with its known effects, including diarrhea, nausea, rash, and hematological events.

Another key difference between masitinib and BTK inhibitors or other drugs developed for MS is that masitinib directly and effectively targets mast cell activity. A growing body of literature highlights mast cells as key contributors to the immunopathology of progressive MS, orchestrating inflammatory responses, BBB disruption, and neurodegenerative processes through complex interactions with glial and immune cells. Therefore, targeting mast cell-mediated pathways offers potential for novel therapeutic strategies to mitigate disease progression and improve clinical outcomes.

Finally, masitinib and BTK inhibitors exhibit distinct safety profiles. Prolonged use of BTK inhibitors, such as zanubrutinib and ibrutinib [FDA 2025; Atallah 2021], has been linked to liver damage, as indicated by elevated liver enzyme levels and, in some instances, drug-induced liver injury, suggesting a potential class effect. Indeed, hepatotoxicity signals have led to clinical holds in MS clinical trials for tolebrutinib, evobrutinib, orelabrutinib and fenebrutinib. In comparison, masitinib has a low risk of hepatotoxicity and a strong long-term safety profile. Considering the overall masitinib safety population, as of February 2024, more than 4,300 patients had received at least one dose of masitinib, including approximately 2,000 patients who received masitinib for over 6 months and 1,200 patients who received it for more than a year [AB Science 2026]. These safety data did not show any correlation between observed asymptomatic elevations in transaminase levels and long-term liver injury. The absence of serious hepatotoxicity, despite the noted increase in enzyme levels, underscores the favorable safety profile of masitinib in MS, suggesting that significant liver toxicity is rare and generally restricted to self-limited, transient liver enzyme elevation without severe clinical consequences.

## Conclusion

Pharmacokinetic data confirmed that orally administered masitinib can achieve clinically relevant concentrations in the CNS, surpassing the inhibitory thresholds necessary for the selective modulation of CSF1R and wild-type Kit, which are crucial drivers of innate immune activation in patients with progressive MS. Alongside indirect indicators of masitinib’s brain penetration (Supplemental Appendix), these findings provide convincing evidence that oral masitinib can remodel the pro-inflammatory microenvironment of the CNS by modulating microglial and mast cell activities. Each of these neuroimmune cells represents a viable target for therapeutic intervention, with the synergistic dual-targeting strategy of masitinib making it a potentially powerful treatment option for progressive MS. Furthermore, this therapeutic approach is possibly well-suited for combination with other treatments that have complementary mechanisms of action and dissimilar safety profiles, such as BTK inhibitors or anti-CD20 monoclonal antibodies, for the treatment of active SPMS or smoldering MS. Masitinib is uniquely positioned to realize the potential of such tailored polytherapy, providing a sound basis for further clinical development across all MS phenotypes, especially progressive forms of MS with the highest unmet medical need.

