## Supplementary Appendix for "Masitinib is an oral, brain penetrant inhibitor of microglial and mast cell activity with neuroprotective potential in progressive forms of multiple sclerosis"

This appendix has been provided to give readers additional information about the study.

- *Supplemental Methods for In Vivo Pharmacokinetic Assessment of Masitinib Brain Penetration in Sprague Dawley Rats*

Given the critical role of innate immune cells, such as microglia, mast cells, and macrophages, in the pathophysiology of progressive multiple sclerosis, it is important to achieve therapeutically relevant central nervous system (CNS) concentrations of masitinib (AB1010) and its active metabolite AB3280 for the effective modulation of these cellular targets. Preclinical data from Sprague Dawley rats provide direct evidence of the ability of masitinib to penetrate the BBB, supporting its potential to engage key immune mechanisms implicated in progressive multiple sclerosis.

Six-week-old male Sprague Dawley rats (Janvier Labs, France) were located in the rodent area of Eurofins ADME Bioanalyses (Vergèze, France) and housed under standard conditions (12-h light/dark cycle) in groups with access to food and water ad libitum. Each animal was identified using an ear tag and examined for general health and welfare. The process, treatment, and euthanasia were conducted according to the relevant procedures used at Eurofins ADME Bioanalyses.

Twelve adult male rats, each weighing approximately 200 g, were administered a single oral dose of masitinib (30 mg/kg). Plasma and brain tissue samples were collected at 2, 4, 8, and 24 h post-administration, with three rats sampled at each time point. Blood samples (0.3 mL per time point) were drawn via retro-orbital sinus puncture under brief isoflurane anesthesia, with precise recording of the sampling times. Following blood collection, the rats underwent intracardial perfusion with 20 mL of saline to clear residual blood from the brain vasculature, ensuring accurate measurement of brain drug concentrations without vascular contamination. This volume is deemed sufficient to clear all blood, given that the blood volume of a rat is approximately 12 mL. The effectiveness of this perfusion was confirmed by the visible blanching of brain tissue. This technique is a terminal procedure because it ends in euthanasia.

Subsequently, brain samples were harvested, homogenized with an Ultra-turrax® in UHQ water (1/1, w/w), and processed through protein precipitation using acetonitrile (100 µl of each homogenate was mixed with 300 µl of acetonitrile, and the mixture was centrifuged for 5 min at 15,000 rpm). The resulting supernatants were analyzed via liquid chromatography-tandem mass spectrometry (LC-MS/MS) using a C18 column to quantify the concentrations of AB1010 and AB3280. Calibration curves ranging from 1 to 2000 ng/mL in brain homogenates and 2 to 4000 ng/g in brain tissue were established, with correlation coefficients exceeding 0.75, ensuring analytical reliability. Plasma samples were centrifuged (2500 rpm) and stored at minus 20°C until analysis, which was conducted using the same LC-MS/MS methodology. This approach provided a robust assessment of the BBB penetrance of masitinib by accurately quantifying AB1010 and AB3280 concentrations in both plasma and brain tissues over time.

**Table S1: AB1010 and AB3280 concentrations measured in plasma and brain after oral administration in male Sprague-Dawley rats (n=3 per time point)**

| Compound | Sampling time (h) | Mean concentration (ng/mL or ng/g) | Standard deviation | Coefficient of variation (%) |
|---|---|---|---|---|
| **AB1010** | 2 | 1121.55 | 663.5 | 59.2 |
| **Plasma (ng/mL)** | 4 | 1492.57 | 454.1 | 30.4 |
| | 8 | 282.75 | 81.0 | 28.7 |
| | 24 | 6.11 | 2.3 | 37.2 |
| **AB1010** | 2 | 159.33 | 77.6 | 48.7 |
| **Brain (ng/g)** | 4 | 223.51 | 55.2 | 24.7 |
| | 8 | 121.53 | 32.9 | 27.1 |
| | 24 | 15.09 | 8.9 | 59.2 |
| **AB3280** | 2 | 53.21 | 33.0 | 62.1 |
| **Plasma (ng/mL)** | 4 | 117.33 | 38.8 | 33.1 |
| | 8 | 21.14 | 5.9 | 27.8 |
| | 24 | 2.81 | 1.5 | 54.8 |
| **AB3280** | 2 | 7.57 | 3.8 | 50.1 |
| **Brain (ng/g)** | 4 | 25.05 | 6.5 | 26.0 |
| | 8 | 33.84 | 8.4 | 24.9 |
| | 24 | 23.72 | 2.10 | 8.86 |

AB1010, masitinib; AB3280, active metabolite of masitinib.
